# Hi-C 2.0: An Optimized Hi-C Procedure for High-Resolution Genome-Wide Mapping of Chromosome Conformation

**DOI:** 10.1101/090001

**Authors:** Houda Belaghzal, Job Dekker, Johan H. Gibcus

## Abstract

Chromosome conformation capture-based methods such as Hi-C have become mainstream techniques for the study of the 3D organization of genomes. These methods convert chromatin interactions reflecting topological chromatin structures into digital information (counts of pair-wise interactions). Here, we describe an updated protocol for Hi-C (Hi-C 2.0) that integrates recent improvements into a single protocol for efficient and high-resolution capture of chromatin interactions. This protocol combines chromatin digestion and frequently cutting enzymes to obtain kilobase (Kb) resolution. It also includes steps to reduce random ligation and the generation of uninformative molecules, such as unligated ends, to improve the amount of valid intra-chromosomal read pairs. This protocol allows for obtaining information on conformational structures such as compartment and TADs, as well as high-resolution conformational features such as DNA loops.

## 1. INTRODUCTION

The spatial organization of chromatin has been a topic of study for many years since chromatin conformation, and long-range associations between genes and distal elements are thought to play important roles in gene expression regulation and other genomic activities. The concept that dense matrices of chromatin interactions could be used to determine the spatial organization of chromatin domains, chromosomes and ultimately entire genomes, was introduced in the original publication that described the chromosome conformation capture method [1]. This concept was tested by development of chromosome conformation capture (3C), its application to yeast chromosomes, and analysis of interaction data using polymer models. This led to the first 3D model of a chromosome.

In 3C, chromatin is first fixed with formaldehyde to covalently link spatially proximal loci. This is essential for efficient detection of chromatin interactions, as leaving out cross-linking leads to dramatic loss of detected contacts and, in our hands, inability to detect chromatin conformation beyond a few Kb. Chromatin is then fragmented with a nuclease and ends are re-ligated. This leads to unique ligation products between spatially proximal loci that can then be detected by PCR, ligation mediated amplification, or direct sequencing.

The concept of using matrices of contact frequencies to infer chromatin folding, and its proof-of-principle in yeast [1] has led to many new studies and the development of a range of 3C-based assays with increased throughput including 4C, 5C and ChIA-PET. Hi-C was introduced in 2009 [2] as a genome-wide version of 3C. The incorporation of biotinylated nucleotides at the digested DNA ends prior to ligation allowed for the specific capture of digested and subsequently ligated chimeric molecules using streptavidin coated beads. These chimeric molecules are then directly sequenced, e.g. on an Illumina platform. Since its introduction, the technique has gone through several stages of adaptation and optimization. We have previously presented a base protocol that was used the incorporation of biotinylated dCTP in an overhang generated by HindIII digestion [3].

Here we present Hi-C 2.0, a further optimized Hi-C protocol that integrates the several technical improvements in a single protocol. One adaptation to the base protocol removes a SDS solubilization step after digestion, which better preserves nuclear structure so that ligation occurs more *in situ*, i.e. in intact nuclei. This prevents random ligation between released chromatin fragments. This adaptation was first introduced for 4C [4] and has since been used for single cell Hi-C [5,6] and more recently for smaller working volumes of Hi-C (*in situ* Hi-C [7]). A second adaptation in recently developed protocols increases the resolution of Hi-C through the use of restriction enzymes that digest more frequently, such as MboI and DpnII, or nucleases such as DNaseI and Micrococcal nuclease [7,9–11]. Thirdly, experimental steps can be included to reduce cost by reducing the number of uninformative sequences such as unligated ends. This is important because even though many topological structures, including compartments and topologically associating domains (TADs) can effectively be resolved by binning 100 million valid pair reads at 100 kb and 40 kb resolution respectively [12–15], detection of point-to-point looping interactions, e.g. between promoters and enhancers or between pairs of CTCF sites typically require >1 billion valid pairs [7]. Therefore, steps to increase the fraction of informative intra-chromosomal reads will help reduce cost.

Alternative approaches to look at looping interactions for specific regions of interest can be captured in a more cost-effective way by targeted approaches such as 4C [16,17], 5C [18,19] and Capture C [20,21].

Here we describe Hi-C 2.0 which uses the DpnII restriction enzyme, in situ ligation, and efficient unligated end removal. A detailed step-by-step protocol is provided in the supplemental materials (“Hi-C 2.0 Protocol”).

## 2. CAPTURING CHROMOSOME CONFORMATION

### 2.1 CELL CULTURE & CROSSLINKING CELLS USING FORMALDEHYDE

The objective to increase the resolution of Hi-C, without dramatically increasing the costs, requires a robust and efficient capturing of spatial DNA interactions, such as between enhancers and promoters. Digesting the genome into more and smaller pieces of DNA increases both the resolution and the complexity of a Hi-C library. To fully capture individual interactions within this complex library of pair-wise interactions, it is helpful to start with a large amount of cells. As such, even very infrequent interactions can still be captured, but in a statistically significant manner. In our Hi-C protocol, we start with 5 Million cells to ensure the generation of complex libraries. Using an adapted version of the protocol described here, we have successfully generated libraries with as little as 500,000 cells of starting material. However, the reduced amount of genome copies made these libraries less complex.

We use a final 1% concentration of formaldehyde to crosslink DNA-DNA interactions that are bridged by proteins. Serum can affect the cross-linking efficiency because it is very rich in proteins and it will compete for formaldehyde. Therefore we replace serum containing medium with serum free medium before fixation.

We distinguish between adherent and suspension cells in order to fix them in their normal growth conditions. Adherent cells are washed once with the relevant serum-free medium before fixation. Fixation occurs by incubation in formaldehyde containing medium without detaching cells from their growth surface. For suspension cells we replace the wash medium with medium containing formaldehyde after centrifugation.

For both cell types, the formaldehyde is quenched with glycine to terminate the crosslinking. Cells are washed with PBS and pelleted cells can be snap-frozen with dry ice or liquid nitrogen. These cells can be stored at −80°C for up to a year before continuing Hi-C.

### 2.2 THE HI- C METHOD

#### 2.2.1 Cells lysis and chromatin digestion

We perform Hi-C on lysed cross-linked cells. We use a douncer to lyse the cells in cold hypotonic buffer that is supplemented with protease inhibitors to maintain Proteins-DNA complexes. After two rounds of douncing we pellet the material and wash twice with a cold buffer that we will use during digestion. At this point an aliquot of ∼ 5% volume can be taken to check the integrity of DNA on an agarose gel.

Before digestion, we incubate the lysed cells in 0.1% of SDS to eliminate proteins that are not cross-linked to DNA, and open the chromatin for a better and more homogenous digestion. The reaction is terminated by addition of triton X-100 to a 1% final concentration. Now the DNA is accessible for digestion by a relevant endonuclease. Previous high resolution Hi-C libraries have used MboI or DpnII [7] to fragment DNA with restriction endonucleases. Alternative ways of digestion include the use of micrococcal nuclease, which digests in between nucleosomes [10] and random breakage by sonication. Here an endonuclease is used that leaves a 5’overhang, which allows marking the sites of digestion with a biotinylated deoxyribonucleotide during overhang fill-in.

Both DpnII and MboI recognize and digest GATC, and leave a 5’-GATC overhang. We prefer the use of DpnII, because it is insensitive to CpG methylation. The GATC sequence is frequently found genome wide and should theoretically result in a median digestion into 256 base pairs fragments for the 3 × 10^9^ base pair (bp) human genome. To ensure maximal digestion, chromatin is incubated with DpnII overnight in a thermocycler with interval agitation. After digestion DNA forms a smear of 400-3000 bp on agarose gel ((Figure 2A-(2). Digestion is terminated by heat inactivation of the restriction enzyme at 65˚C for 20 minutes.

#### 2.2.2. Marking of DNA ends with biotin

DNA digestion generates a 5’overhang that is then filled in with deoxyribonucleotides. By strategically replacing one of the deoxyribonucleotides with a biotin-conjugated variant, we can mark the site of digestion and enable enrichment for those sites in a later step. It is this specific fill-in that separates Hi-C from other chromosome conformation capture based methods. For DpnII, we incorporate biotin-14-dATP (Figure 1C). Although the incorporation of biotinylated dCTP is theoretically possible, we have found that this incorporation of a biotinylated nucleotide at the end the overhang leads to less efficient ligation (below).

**Figure 1.**
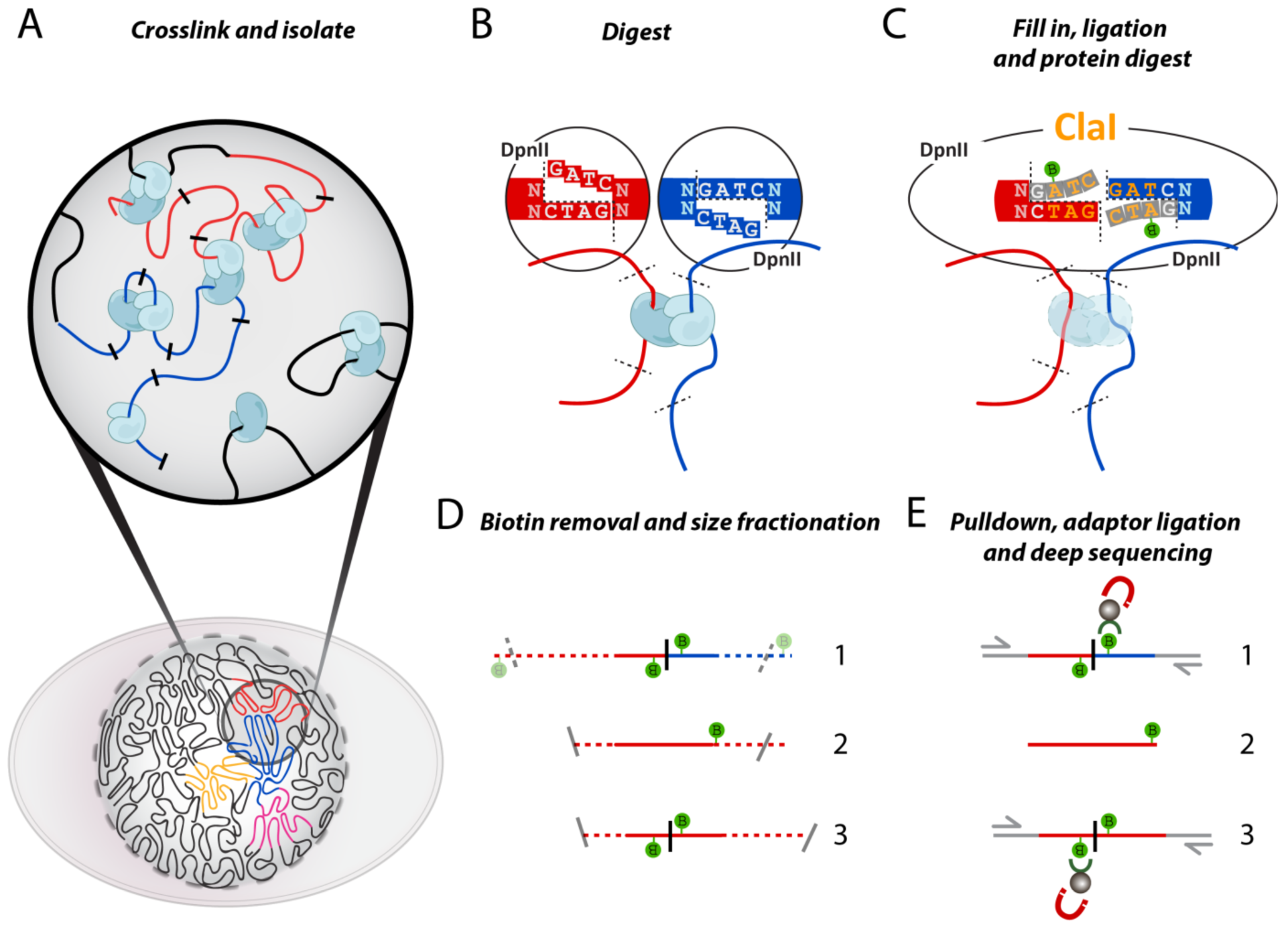
Overview of the Hi-C method. **(A)** Cells fixed with formaldehyde contain protein-mediated DNA-DNA interactions. **(B)** DNA digestion with DpnII, recognizing GATC and generates a 5’-GATC overhang. **(C)** Filling in of the 5’overhang with dNTPs and biotin-14-dATP blunts the overhang. Ligation of the blunted ends creates a new restriction site (ClaI), which can be used to assess fill-in efficiency. After ligation, crosslinks are reversed to remove proteins from DNA. **(D)** Removal of Biotin from un-ligated ends. DNA is fragmented to 200-300bp DNA fragments to enable paired-end sequencing. **(E)** Enrichment of ligation junctions by using the high affinity of streptavidin coated beads for the incorporated biotin allows for ligation product enrichment prior to adapter ligation.

Klenow fragment of DNA polymerase I is used to fill in the 5’ overhang for 4 hours at 23°C. This low temperature is crucial for efficient incorporation of the large biotinylated dATP and decreases 3’  5’exonuclease activity. Not all overhangs will be filled to completion; by consequence not all digested fragments can be properly ligated. In a later step, after DNA purification, unligated biotinylated ends are removed to ensure that only proper ligations are captured and sequenced.

#### 2.2.3. In situ Ligation of proximal ends

Before starting ligation a 10 μl aliquot is taken that will be used to assess digestion efficiency on an agarose gel (Figure 2A-2). The size of the digested DNA is then compared to DNA that was kept aside after lysis before digestion and the DNA that is to be isolated from our ligated Hi-C library. While previous protocols used SDS to inactive the restriction enzyme prior to ligation, here we describe the “*in situ*“ ligation protocol [4,5,7], which leaves out this step and inactivates the restriction enzyme by heat. Leaving out this SDS step better preserves nuclear structure and reduces random ligation. Chromatin is then ligated for 4 hours at 16°C, which in our hands is efficient for most of Hi-C libraries. However, in some cases increasing the ligation time to improve the ligation efficiency may be needed. Note that prolonged ligation may increase random ligation. Ligation of the 2 blunted ends creates a new restriction site that be used to assess the ligation efficacy (Figure 1C).

**Figure 2.**
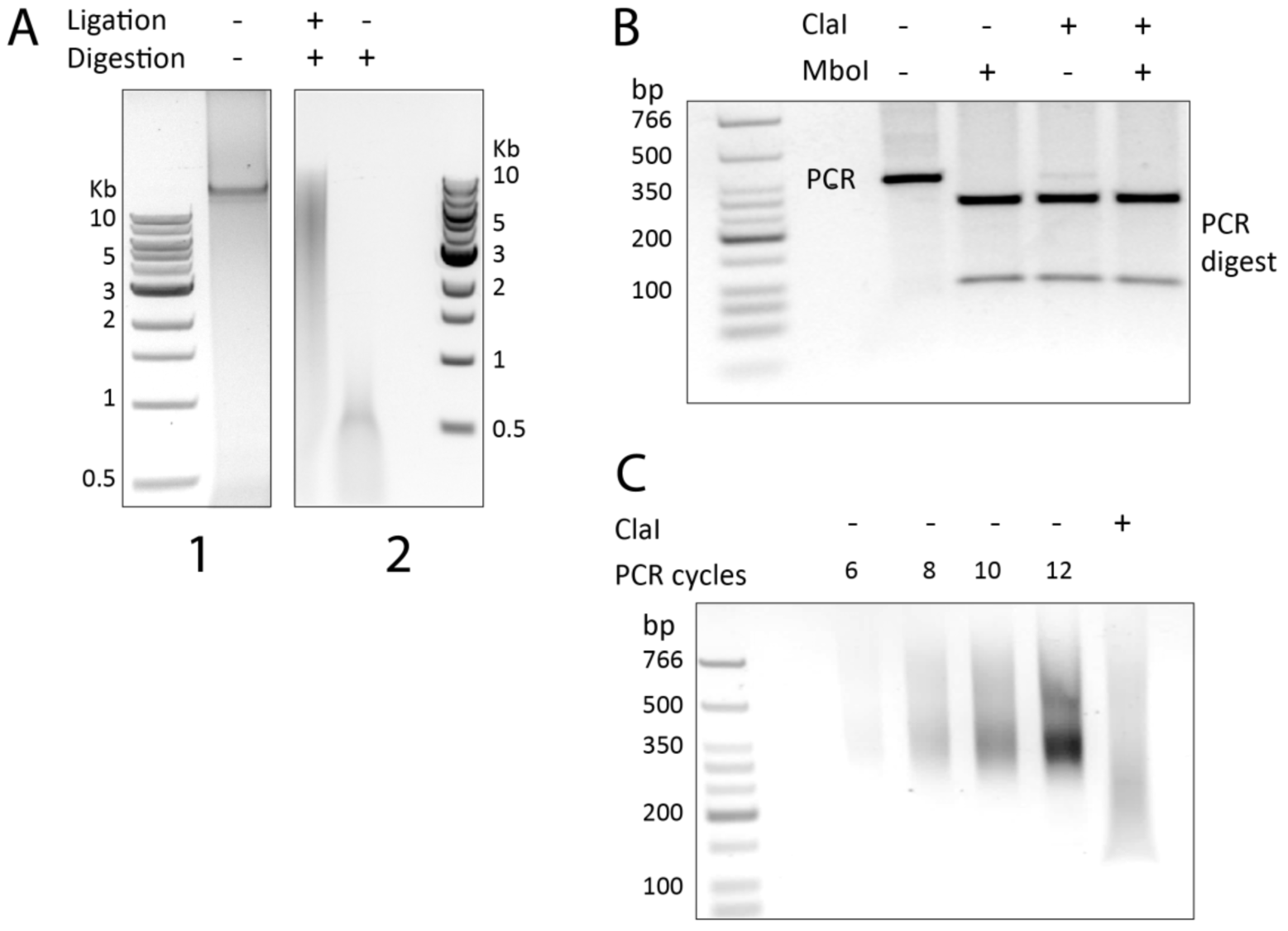
Quality Control of Hi-C ligation products. **(A.1)** Quality control of intact genomic DNA after cell lysis and before digestion. (A.2) Hi-C DNA after digestion and ligation (+,+) compared to unligated, digested control (-,+). Size is indicated by the 1Kb Molecular Weight Ladder from NEB (1 and 2). **(B)** PCR amplification of a specific ligation product to assess ligation efficiency. The PCR product (lane 1), PCR product digested with MboI (lane2), ClaI (lane3), or both ClaI and MobI (lane4). Only properly filled-in ligation products will be digested with ClaI. This allows for a qualitative comparison to MboI digestion, which cuts GATC sites that are present at the ligation junction of both properly filled-in and non-filled-in ligation products. Digestion of the PCR product using ClaI indicates efficient fill-in. The molecular weight ladder used is the Low Molecular Weight Ladder from NEB. **(C)** PCR titration of the final Hi-C library and quantification of the fill-in and ligation efficiency by ClaI digestion. PCR amplification is performed with primers that recognize the PE adaptors that were ligated to the Hi-C library before sequencing. With 6-cycles of PCR amplification enough DNA was produced for sequencing (lane #1). The last lane shows a downward shift of the amplified library after digestion with ClaI, indicative of efficient fill-in.

This blunt end ligation can lead to specific chimeric ligations between ends that were in close proximity after crosslinking. However, this process can also generate circularized ligation products of single restriction fragments. These are not informative and are not considered valid pairs (Figure 3B-(3).

#### 2.2.4. Reversal of crosslinking and DNA purification

Now that interacting loci are ligated into chimeric pieces of DNA, proteins that hold interacting fragments in close proximity can be removed. This is achieved by thermal reversion of crosslinked proteins and incubation with uproteinase-K.

After proteinase K treatment DNA is isolated using 2 steps of phenol:chloroform (pH=7.9) and DNA is precipitated using a standard sodium acetate plus ethanol protocol. An Amicon column is used to wash pelleted DNA with low EDTA, tris-buffered water (TLE) to remove any excess of salt.

#### 2.2.5. Quality Control of Hi- C ligation products

During the procedure described above, small aliquots were taken after three key steps in the protocol: lysis, digestion and ligation. DNA isolated from these aliquots can be run on an agarose gel to ascertain the intactness of the DNA prior to digestion, the extent of digestion and efficiency of subsequent ligation. The undigested genomic DNA typically runs as a tight band of over 20 Kb in size (Figure 2A-(1). After digestion, the DNA runs as a smear with a size range specific for the applied restriction enzyme (Figure 2A-(2). Both of these controls allow for a comparison with the actual library of DNA containing the chimeric ligated ends. These ligated chimeras should have a higher molecular weight than the digestion control and are most likely smaller in size than the undigested control. For DpnII digestion we usually obtain sizes ranging between 3Kb and 10kb (Figure 2A-(2).

A second quality control involves quantification of the level of fill-in of overhangs prior to ligation. This is done by PCR amplification of a specific ligation product with primer pairs that were designed for 2 nearby digestion sites (e.g. adjacent restriction fragments) followed by digestion of the PCR product with a restriction enzyme that only cuts at the ligation junction when fill-in has occurred prior to ligation.

Specifically, PCR reactions are set up to detect head-to-head ligation products (Figure 3B and (C). Primers are designed near neighboring restriction sites that have a high likelihood of being in close spatial proximity, which can only generate PCR products when properly ligated chimeras are present (Figure 3B). For some endonucleases, including HindIII and DpnII, ligation of the 2 blunt ends generates a new digestion site that can be used to quantify the ligation efficiency (Figure 1C). After PCR amplification of a ligation product, the PCR product is digested with the enzyme that recognizes this newly generated ligation product (Figure 2B). Typically the majority of the PCR product is cleaved indicating efficient fill in.

### 2.3 PREPARING CAPTURED CONFORMATIONS FOR DEEP SEQUENCING

#### 2.3.1. Removal of Biotin from un-ligated ends

We have found that in most Hi-C experiments some digested sites will have remained unligated. For example, if the fill-in of some overhangs was incomplete, ligation to a proximal fragment will not occur and the overall ligation will not be 100% efficient. Such cases result in biotinylated but unligated ends. We prefer to remove these “dangling” ends from the Hi-C library, because they would make sequencing less efficient by generating uninformative reads (Figure 1D).

Our biotin removal step uses T4 DNA polymerase and a low concentration of dNTPs to favor the 3’ to 5’ exonuclease activity over its 5’ to 3’ polymerase activity. By only providing dATP and dGTP, which are complementary to the inside of the 5’ overhang, the polymerase will not be able to complete re-filling the overhang after removing the filled in bases.

Importantly, this biotin removal step significantly reduces the amount and variability of dangling ends to just 1-2%. In our hands, without this step the dangling end frequency is quite variable between experiments and can be as high as 30%. Therefore the relative increase in valid pairs of DNA sequences will effectively generate more relevant reads while not increasing cost.

#### 2.3.2. Sonication

In order to sequence both ends of ligation products DNA is sonicated to reduce their size to 200-300 bp in preparation for paired-end sequencing. For sequenced reads to be mapped correctly, each end of a paired-end read should not pass the chimeric ligation junction, since this will result in a sequence that cannot be identified in a reference genome. Fragements that are 200-300 bp are likely to contain enough mappable sequence at each end before reaching a ligation junction. We prefer to use a Covaris sonicator, because is highly reliability and reproducible in generating a tight range of DNA fragments.

#### 2.3.3. Size selection

Covaris sonication results in a relatively small size range of DNA fragments. Therefore, additional size selection can be omitted, but we prefer to use SPRI beads (AMpure) to create an even tighter distribution of fragments. Ampure is a mixture of magnetic beads and polyethylene glycol (PEG-8000). Adding AMpure to a DNA solution reduces the solubility of DNA, because PEG, a crowding agent, will effectively occupy the hydrogen bonds of aqueous solutions. As a result of this crowding, DNA will come out of the solution and bind to the coated magnetic beads. Since larger DNA molecules will come out of solution first, the final concentration of PEG can be used to generate a size cut-off. After sonication 2 consecutive size selections with Ampure are performed. The first AMpure selection will precipitate DNA larger than 300 bp. Using a magnet, bead-bound DNA is separated from the PEG supernatant, which contains fragments smaller than 300 bp. This supernatant undergoes an additional AMpure selection that precipitates DNA larger than 150 bp. Here after, the bead-bound DNA will be narrowly sized to 150-300 base pairs.

#### 2.3.4. End repair

The shearing of DNA by sonication will inevitably damage DNA ends. To repair all the ends after sonication a mix of T4 and Klenow DNA polymerase is used together with T4 polynucleotide kinase (PNK). The first 2 enzymes will repair nicked DNA and single stranded ends, while T4 PNK phosphorylates 5’-ends allowing subsequent A-tailing and adaptor ligation.

#### 2.3.5. Biotin pulldown

To enrich for Hi-C ligation junctions, we use streptavidin coated beads with a high affinity for the incorporated biotin. This effectively eliminates any DNA without biotin, i.e. DNA that wasn’t properly digested, filled-in and ligated (Figure 1E).

As mentioned above, the step to remove biotin at DNA ends (see 2.3.1) reduces the pulldown of a large fraction of unwanted unligated fragments. However, some unwanted fragments might still be captured. These include self-circled ligation products and other fragments that were insensitive to biotin removal (Figure 3B-2 and B-3). For instance, during biotin incorporation, internally nicked DNA could be repaired with biotinylated nucleotides and when too far away from the DNA ends, these incorporated biotinylated nucleotides will not be removed by T4 Polymerase in our biotin removal step. These read pairs will need to be removed during bioinformatic analysis of the data.

#### 2.3.6. A-tailing and adaptor ligation

For DNA sequencing Illumina PE adaptors are ligated to both ends of the size selected ligation products. The PE adaptors were generated from DNA oligos, which after duplexing have a 5’-dTTP overhang. This overhang increases ligation efficiency when presented with a free 3’Adenyl. The 3’end of the ligation products are adenylated using dATP and a Klenow fragment lacking 3’ to 5’ exonuclease activity, and then adaptors are ligated using T4 DNA ligase. Depending on whether the preferred sequencing protocol includes multiplexing, one can either use indexed or non-indexed paired-end adapters. We have successfully used paired-end single index adaptors to sequence multiple libraries in a single lane. Strategically choosing the right combination of multiplex adaptors, as suggested by Illumina, at this step is essential when multiplexing is intended.

#### 2.3.7. PCR titration and production

To obtain enough DNA for deep sequencing, the library of ligated fragments is amplified by PCR, using primers designed to anneal to the PE adaptors. Since over-amplification by PCR can result in reduced library complexity, a PCR titration is performed on an aliquot to find the optimal amount of PCR cycles. The smallest number of PCR cycles, producing enough DNA for sequencing will be chosen (Figure. 2C). After PCR, the PCR product re separated from the bead-bound DNA for a final AMpure cleanup.

As a final quality control an aliquot of the amplified library is digested with ClaI (Figure 2C).Ligation of blunted DpnII sites creates a new ClaI site at the ligation junction. When fill in and ligation was successful and efficient one expects the majority of the PCR products to be cleaved, resulting in a shift in size compared to undigested PCR product which can be observed by running an aliquot of DNA on an agarose gel (Figure 2C).

### 2.4. Sequencing

The Hi-C library is then sequenced. We generally sequence these fragments using Illumina 50 bp paired-end sequencing. Using longer paired end reads (e.g. 100 bp instead of 50 bp) can increase the number of reads that can be uniquely mapped. In our hands, using 100 bp paired-end reads increases the total number of read pairs where both ends can be uniquely mapped by as much as 15-20 % as compared to using 50 bp paired end reads.

## 3. DATA ANALYSIS

### 3.1 MAPPING AND BINNING PIPELINE

The paired sequencing information can be downloaded from the sequencing platform as standard fastq files. The reads are then mapped to a reference genome and valid interaction pairs are identified, while uninformative reads like self-circles and dangling ends are removed (Figure 3). Valid interaction pairs can be binned at a range of resolutions (e.g. 5-100 Kb bins) [22]. There are several public pipelines for processing Hi-C data, e.g. HOMER [23], HiCPro [24] and Juicer [25]. For a more detailed description of mapping and binning of data using our pipeline, we refer to in Lajoie et al., *Methods* (2015) [22]. A high quality Hi-C library for mammalian genomes typically has >70% of interactions mapping to intra-chromosomal interactions, less that 1-4 % unligated ends, less than 1-2% self ligated circles, and less than 5% PCR redundant interactions per 400 million reads. We note that in our experience some sequencing platforms can produce additional apparently redundant reads on the flow cell that are not due to PCR amplification but to loading of the sample. Finally, these numbers can depend on biological state and therefore are only general guidelines for assessment of library quality.

**Figure 3.**
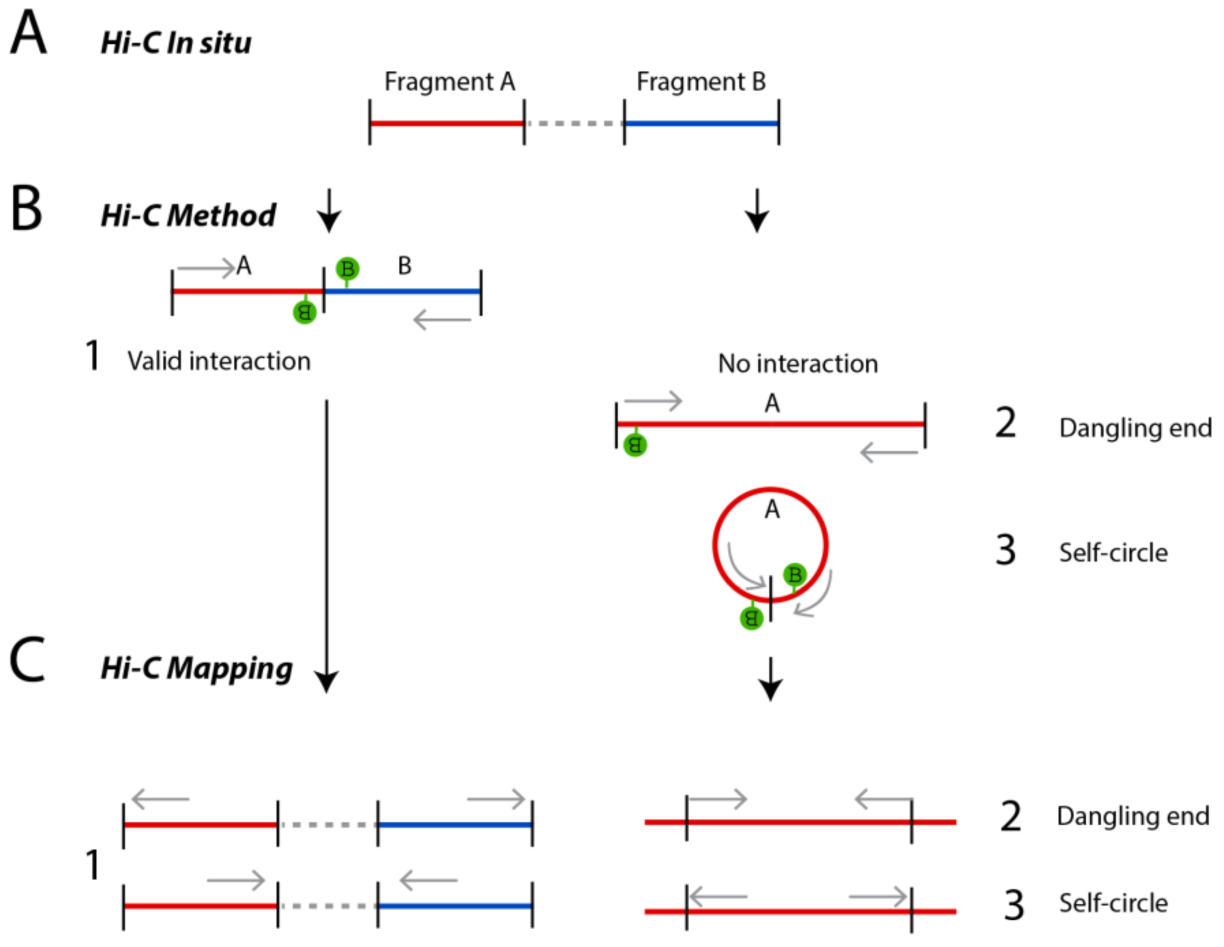
Possible products generated with Hi-C. **(A)** Fragment A and B are not neighboring in the linear genome. **(B)** If fragment A and B are in close spatial proximity they can become cross-linked and ligated during the Hi-C procedure (1). Other possible non-valid products can be derived from non-ligated DNA (dangling-end; 2) or single fragments that have become circularized after ligation (self-circles; 3). The gray arrow indicate the orientation of the paired-end reads in the Hi-C library **(C)** Dangling ends can be removed from the Hi-C library prior to sequencing, as described in this protocol. Any remaining dangling-ends and self-circles can be filtered out from the sequenced library computationally after mapping and assessing the orientation of the DNA. After mapping, valid reads locate to different fragments in the reference genome and are either inward or outward oriented. Invalid reads map to the same fragment in the reference genome and can also be either inward (dangling ends) or outward oriented (self-circles). Gray arrows indicate the read orientation in the reference genome for both valid pair chimeras (1) and non-valid pair interactions such as dangling-ends (2) and self-circles (3)).

Binned reads are stored as a symmetric matrix with each row and column representing a genomic location (bin). Interacting regions are represented by the number of reads for every bin within this matrix. These matrices are routinely displayed as heatmaps that display these interactions by coloring bins for the amount of reads they contains. The diagonal within these maps (x=y) represents neighboring interactions.

### 3.2 OBTAINING TOPOLOGICAL INFORMATION

Recent research has shown that the genome is composed of several layers of structure, ranging from compartments to topologically associating domains (TADs) and loops (Figure 4). We will briefly describe how those features can be measured. For more details we refer to Lajoie et al., *Methods* (2015) [22].

**Figure 4.**
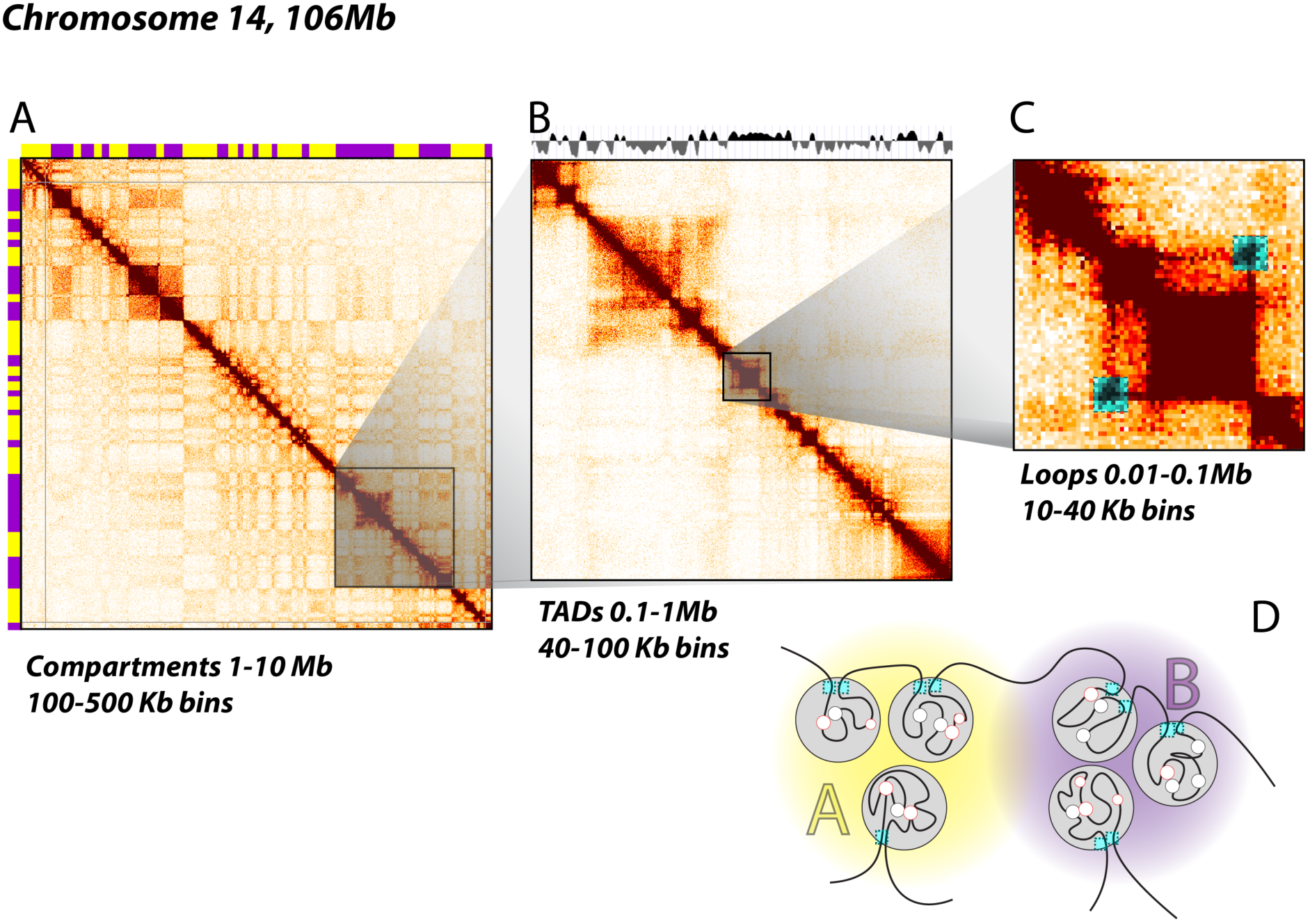
Topological structures obtained at increasing resolution. **(A)** Heatmaps generated from 100 kb binned Hi-C data for chromosome 14 show the alternating pattern of A and B compartments (yellow/purple) **(B)** On a sub-chromosomal level, heatmaps at 40 kb resolution show the location of TADs, as indicated by an insulation score on top (gray). **(C)** Within TADs, DNA loops can form that show up as “dots” of interactions in heatmaps of sufficient resolution (typically 10 Kb bins or less). **(D)** Interpretation of the topological hierarchy obtained from Hi-C. TADs (gray circles) within the same compartment (A or B) interact more frequently than between different compartments. TADs are bordered by insulating proteins (e.g. CTCF, cyan squares). DNA loops form between CTCF sites, enhancers and promoters (red/black circles).

#### 3.2.1. Compartments

Compartments are defined as groups of domains, located along the same chromosome or on different chromosomes that display increased interactions with each other. In heatmaps generated from 100Kb bins, this is visible as a specific plaid pattern. These alternating blocks of high and low interaction frequencies represent A and B compartments [2]. Principal component analysis (PCA) readily identifies these compartments that tend to be captured by the first component. The active “A” compartments are gene-dense euchromatin regions, whereas the inactive “B”-compartments are gene-poor heterochromatin regions (Figure 4).

#### 3.2.2. Topologically associated domains (TADs)

TADs are contiguous region that display high levels of self-association and that are separated from adjacent regions by sharp boundaries[13,15]. The locations of TADs can be determined when interaction data is binned at 40 Kb or less. There are several computational approaches to identify the locations of TAD boundaries including the directionality index [13] or an insulation score algorithm [26] to determine the location of TADs (Figure 4).

#### 3.2.3. Point to point interactions (loops)

Many point-to-point interactions or loops appear as off-diagonal “dots” in a heatmap. Typically, a 10Kb resolution or higher is required for visualizing looping interactions. Mapping to smaller bins will allow for more specific interactions, but this comes at the cost of a decreased number of reads per bin. Specific interactions between for instance pairs of CTCF sites are expected to show up as increased signal compared to their surrounding area[7]. Rao et al describe a useful approach to detect such dots using a local background model (Figure 4) [7]. Other types of local interactions, e.g. lines in the heatmap can be detected using global background models [27,28].

## 4. CONCLUSIONS

This Hi-C 2.0 protocol combines in situ ligation with dangling end removal to produce Hi-C libraries enriched in intra-chromosomal valid interaction pairs. This protocol can effectively be used to visualize chromosome conformation at Kb resolution genome-wide.

## ACKNOWLEDGEMENT

We would like to thank Liyan Yang (Dekker lab) and Sarah Hainer (Fazzio lab) for their help in developing this protocol. Work in the Dekker lab is supported by the National Human Genome Research Institute (R01 HG003143, U54 HG007010, U01 HG007910), the National Cancer Institute (U54 CA193419), the NIH Common Fund (U54 DK107980, U01 DA 040588), the National Institute of General Medical Sciences (R01 GM 112720), and the National Institute of Allergy and Infectious Diseases (U01 R01 AI 117839). The authors declare that they have no competing interests. J.D. is an investigator of the Howard Hughes Medical Institute.

